# Social ageing can protect against infectious disease in a group-living primate

**DOI:** 10.1101/2024.03.09.584237

**Authors:** Erin R. Siracusa, Melissa A. Pavez-Fox, Josué E. Negron-Del Valle, Daniel Phillips, Michael L. Platt, Noah Snyder-Mackler, James P. Higham, Lauren J. N. Brent, Matthew J. Silk

## Abstract

The benefits of social living are well established, but sociality also comes with costs, including infectious disease risk. This cost-benefit ratio of sociality is expected to change across individuals’ lifespans, which may drive changes in social behaviour with age. To explore this idea, we combine data from a group-living primate for which social ageing has been described with epidemiological models to show that having lower social connectedness when older can protect against the costs of a hypothetical, directly transmitted endemic pathogen. Assuming no age differences in epidemiological characteristics (susceptibility to, severity, and duration of infection), older individuals suffered lower infection costs, which was explained largely because they were less connected in their social networks than younger individuals. This benefit of ‘social ageing’ depended on epidemiological characteristics and was greatest when infection severity increased with age. When infection duration increased with age, social ageing was beneficial only when pathogen transmissibility was low. Older individuals benefited most from having a lower frequency of interactions (strength) and network embeddedness (closeness) and benefited less from having fewer social partners (degree). Our study provides a first examination of the epidemiology of social ageing, demonstrating the potential for pathogens to influence evolutionary dynamics of social ageing in natural populations.

## Introduction

Social interactions have an important influence on individual fitness through their effects on survival [1] and reproductive success [2–5]. However, sociality is not exclusively beneficial but comes with important costs ranging from competition for food, mates, and other resources [6–8] to increased risk of pathogen and parasite transmission [9,10]. Understanding how these costs and benefits are balanced is central to understanding the evolution of social living [11].

Recent research indicates that social interactions and relationships are not stable over time but differ across the lifespan [12–15]. There are multiple hypotheses for why individuals might change their social behaviour as they get older, including physical and cognitive declines [16,17], demographic changes and shifting kinship dynamics [18,19], or enhanced skill and experience with age [20]. Fundamentally, the costs and benefits of interacting with others may change across the lifespan [12], leading individuals to actively adjust their social behaviour in response. Recent work, both in humans and non-human animals, suggests older individuals tend to have fewer affiliative social partners than their younger counterparts [13,15,21–23], spend less time in affiliative interactions [13,22–24], and to be less well embedded in the wider social network [23,25,26]. There is evidence to suggest that in some species, at least, this is the result of older individuals being more selective with whom they interact (i.e. ‘social selectivity’ [21,27,28]). This evidence is consistent with the hypothesis that age-based reductions in social connectedness (i.e. ‘social ageing’) are not exclusively the by-product of declines in bodily systems but instead reflect beneficial behavioral changes that ameliorate senescence-associated costs [21].

One major cost of socialising, which has the potential to be exacerbated in old age, is the increased risk and costs of contracting infectious diseases [29], with individuals that interact more with others at greater risk of infection [30,31]. Therefore, one appealing hypothesis is that, because immunosenescence often means that individuals are less able to fight infections as they age and so suffer greater morbidity to infectious diseases [32–34], age-based reductions in individual social network connectedness can help mitigate disease costs in older individuals [35]. By being socially selective (i.e., reducing their number of social partners while socialising for longer with their closest associates [21,27]), older individuals may be able to reduce their risk of infection while maintaining the benefits of social relationships [36,37]. However, we currently lack evidence that these declines in sociality with age (hereafter referred to as social ageing) are sufficient to protect an individual from infectious disease.

One challenge with demonstrating the disease-associated consequences of social ageing is the fact that an individual’s social interactions and relationships are embedded in a wider group or population wide social network. These social networks are complex, emergent structures that depend in a non-linear way on the behaviour of every individual [38]. As a result, it can be difficult to predict both the consequences of individual behaviour change on the group-level social structure and the knock-on implications for pathogen spread across the network. While connectivity (or social network density) is an important component of the vulnerability of a group or population to an infectious disease outbreak [10], other aspects of social network structure such as the transitivity of social connections (the tendency of an individual’s connections to also be connected to each other) and modularity (division of the network into a set of social communities) can also shape how pathogens spread [9]. Consequently, testing whether being more socially selective with age can protect older individuals from infectious disease costs requires the use of epidemiological network models [39,40].

Rhesus macaques (*Macaca mulatta)* offer an excellent system in which to test this question. They live in matrilineal multi-male multi-female social groups [41] where the social relationships that individuals form have important effects on their fitness [8,20,42,43]. A long-term study of a free-ranging population of rhesus macaques on Cayo Santiago Island, Puerto Rico, has generated a unique long-term dataset encompassing multiple social groups that has provided important insights into primate social behaviour [44]. Recent research investigating patterns of social ageing in female rhesus macaques has demonstrated that as females get older they reduce their number of social connections and spend more time socializing with important partners (such as kin and partners with strong and stable connections; [21]). These changes resulted from females approaching fewer partners as they got older, while continuing to spend the same amount of time socializing, and were not driven by partners dying or avoiding them.

This study provides one of the few examples in non-human animals where active reductions in sociality with age have been distinguished from more passive processes such as declines in social engagement driven by the loss of social partners, declining social motivation, or impaired physical ability to engage [21]. The results suggest that age-based reductions in an individual’s number of social partners might enable individuals to cope with the physiological alterations or limitations that come with age, such as immunosenescence. Female macaques also show changes in their indirect connectedness (connections to partners of their social partners) with age, with measures of both betweenness (bridging capacity) and closeness (ability to reach others or be reached in the network) declining as females get older [25]. In addition to showing clear patterns of social ageing, rhesus macaques are a powerful system to explore the intersection of social ageing and disease risk because they are popular biomedical models of human health and ageing [45–47] and are frequently used as a model species in studies of infection biology, thereby providing information on the types of pathogen they are susceptible to [45] and how costs of infection vary with age [48–51]. The fact that the observed patterns of social ageing in macaques closely resemble those observed in human populations [21], combined with the similarity between macaques and humans in patterns of immunosenescence [45,47,52] and susceptibility to pathogens (such as SARS-CoV-2 [53,54]) makes this a uniquely relevant system for understanding the potential implications of age-based changes in sociality for disease risk in humans.

Here we combined epidemiological network models of pathogen spread with empirical data on social ageing from a population of free-ranging rhesus macaques. Using susceptible-infected-susceptible (SIS) models of the spread of a hypothetical directly-transmitted pathogen (e.g. respiratory virus) across 23 real-world rhesus macaque social networks, we hypothesised that lower social connectivity of older individuals [21] would protect them from accumulated infection costs of an endemic (constantly present) pathogen. Specifically, we quantified how age-based variation in three measures of social centrality - degree (number of partners), strength (amount of time spent socializing) and closeness (capacity to reach or be reached in the network) influenced infection costs with age. Our choice of social metrics was based on prior knowledge that individuals in this system show within-individual declines in both degree and closeness with age, but not in strength [21,25]. We explored the effects of strength given it is a measure of social connectivity that is well-known to have important consequences for disease transmission [30]. We assessed how an individual’s accumulated costs of infection in each simulation were influenced by their age and the aforementioned network measures of social connection.

We predicted that, (1) across all measures of social centrality, the cost of infection would decrease with reduced social connectedness in an age-dependent manner, such that for older individuals it would be much less costly to have lower connectedness in the social network compared with younger individuals where the benefits of having lower connectedness would be more moderate. Next, we predicted that (2) most of the decrease in infection cost associated with social ageing would be driven by older individuals having lower degree and closeness since strength was not expected to change with age in this population. We further predicted that (3) social ageing would provide the greatest benefit, in terms of reducing infection cost, when immunosenescence, and therefore infection cost with age, was greatest. To assess this we explored the extent to which these social variables influenced infection cost across a range of pathogen transmissibilities and under conditions where we varied different components of immunosenescence, including how susceptibility to (likelihood of acquiring infection), as well as severity (per timestep cost of infection) and duration of infection changed with age.

For the sake of clarity and brevity, throughout the manuscript we refer to this reduction in infection cost that is the result of older individuals having lower social connectivity as the ‘protective effect of social ageing’ although our analyses are at the population level rather than being exclusively within-individual, and this interpretation therefore warrants caution. However, given that our previous results [21,25] have shown that age-based differences in sociality are driven by within-individual changes rather than between individual differences (at least for degree and closeness), it is appropriate to assume that population level trends are the result of within-individual processes [55,56]. Our models are the first to explicitly test whether reduced social connectivity among older individuals is sufficient to buffer against infectious disease costs in a free-ranging population.

## Methods

### Study System

We studied a population of rhesus macaques on the island of Cayo Santiago off the southeast coast of Puerto Rico. This population was first introduced to the island in 1938 from India and currently consists of ∼ 1800 individuals living in 12 mixed-sex social groups. The population is maintained by the Caribbean Primate Research Center, which is responsible for monitoring the population daily and collecting data on births, deaths, and social group membership. The animals are food supplemented and provided *ad libitum* access to fresh water. The island is predator free and there is no veterinary intervention for sick or wounded animals, meaning that the majority of deaths on the island are from natural causes such as illness and injury [8].

For this study we focused on adult female rhesus macaques (aged 6 years and older) from six social groups that have been studied intensively between 2010 and 2022 and therefore for which we had detailed behavioural data to build social networks. In total we used behavioural data collected from twenty-three different group years (group F 2010-2017, group HH 2014 & 2016, group KK 2013, 2015 & 2017, group R 2015 & 2016, group S 2011 & 2019, group V 2015-2017, 2019, 2021-2022). For these analyses we excluded data collected in 2018 and 2020 because Hurricane Maria and the COVID-19 pandemic, respectively, precluded use of our typical protocol to collect behavioural data. In total this resulted in 1176 macaque-years of data from 410 unique females whose ages ranged from 6-28 years old (mean = 11.2; see Fig. S1). We collected behavioural data between 07:30 - 14:00, which are the working hours of the field station. Behavioural data were collected using 10-min (20 group years) or 5-min (3 group years) focal animal samples, where all behaviours were recorded continuously. We stratified sampling to ensure balanced data collection on individuals throughout the day and over the course of the year. Given that previous research in this system has shown that there are clear age-based changes in grooming associations among female macaques [21,25], in addition to the fact that behaviours with prolonged contact (such as grooming) are known to be highly relevant for parasite transmission [57–62], we focused specifically on grooming interactions to build our social networks (see below). During focal observations, we recorded the duration of the grooming bout as well as the identity of the monkeys and the direction of the grooming behaviour (give or receive). Grooming bouts had to be at least 5 seconds long to be recorded and a new bout of grooming was recorded if the identity of the monkeys or direction of grooming changed or there was at least a 15 second pause in grooming behaviour. These thresholds are based on long-term expert knowledge of the study system and have been used since 2010 in the collection of grooming interactions. We focused our study on adult females because the age-based changes in sociality previously demonstrated in this system were for female networks that excluded interactions with males and juveniles or subadults of either sex [21,25]. Focusing on female-female interactions also allowed us to isolate how changes in cooperative interactions with age influence disease transmission outside of age-based changes in socio-sexual behaviour. This is because females’ interactions with males are likely to capture both cooperative interactions as well as reproductive behaviours, making it difficult to parse age-based changes in cooperation from age-based changes in reproduction.

### Social network construction

We constructed grooming networks for 23 group years (6 groups across 12 years; mean = 3.8, range = 2-8 years per group). In these networks, nodes represent individuals and edges represent the undirected rate of grooming between a pair of individuals (number of grooming bouts/total number of focals of both individuals). Although we have data on both grooming given and received, here grooming serves as a proxy for time spent in close contact. We expected that the total amount of time and number of partners with which an individual spent time in close contact was likely to be most relevant for infection risk from a hypothetical directly-transmissible pathogen, regardless of directionality. Because our social networks were constructed from observational data, which represent a sample of the interactions between individuals, we used the R package bisonR [63] to estimate uncertainty in the quantified edge weights based on sampling effort. Explicitly incorporating uncertainty around the observed social network allows us to account for how well the estimated network represents the true underlying latent network and therefore confirm the robustness of our modelling results to sampling effects. For example, typical network approaches would assign an edge weight of 0.5 both for a dyad that had been seen together once and apart once as well for a dyad that had been seen together 100 times and apart 100 times, despite the fact that our certainty about the edge weight of the second dyad is much greater than the first [64]. By generating a distribution of possible networks from the observed data rather than a single network, BISoN (Bayesian Inference of Social Networks) allows us to account for this uncertainty, which can drastically affect the performance of statistical models [64]. Specifically, for each group year we fitted a Bayesian ‘edge model’ with a count conjugate prior to our observed network data, which returns a posterior distribution of edge weights for each dyad in our network rather than a point estimate (for more information see [64,65]). We extracted 1000 draws from this posterior distribution and used these draws as the social networks over which we modelled pathogen spread. From each BISoN network we calculated three different social centrality measures for each individual: strength (weighted sum of an individual’s social connections); closeness (the inverse of the mean weighted path length from each individual to all others in the network); and degree centrality (the number of social connections each individual has). All three measures were chosen because they have well-established and important consequences for pathogen transmission [30,66–68], and degree and closeness are known to decline with age in this system [25,69]. BISoN model outputs include only non-zero edge weights because even though a particular dyad may never have been seen interacting, BISoN naturally accounts for the possibility that those individuals may have interacted in a future sample and so computes uncertainty for all edge weights [64]. Therefore, in order to calculate degree centrality, we set a threshold at which individuals were deemed to have a social connection or not. As females age, they lose their weakest social connections first. To best capture this change, and ensure that our measure of degree was as independent of strength as possible, we set our threshold using the minimum empirical observed non-zero edge weight for each group-year (mean = 0.017; range = 0.008 - 0.027). We therefore calculated degree centrality by counting all edges in each BISoN network that were equal to or greater than this minimum edge weight in the observed network for each group-year.

### Epidemiological model

#### Overview

We modelled SIS (susceptible-infected-susceptible) epidemiological dynamics to simulate the spread of a hypothetical, directly-transmitted endemic pathogen by close contact through our social networks (e.g., a respiratory virus). In our modelling framework individuals could either be susceptible (S) or infected (I), thus we assumed that individuals retained no immunity from previous infections. In a baseline (‘control’) version of the model, the probability of infection depended on interaction strength in the grooming network (parameter: *si*; more grooming = higher transmission probability), individuals were then infected for 5 time-steps (parameter: *di*) and accumulated 1 unit cost per time step that they were infected (i.e., baseline per time step cost - parameter: *ci*). While our per time step cost value was arbitrary, we were interested in the relative differences between individuals rather than the absolute values of this parameter as our goal was to compare infection cost between individuals of different ages and social centralities. As a result, the exact value of this cost parameter was not important. In total, the model ran for 500 time-steps allowing individuals to be infected multiple times within each simulation. We quantified their total cost of infection during a simulation. We then ran additional iterations of the model where we allowed for immunosenescence in three different individual characteristics (infection susceptibility, infection duration and infection severity) based on previous evidence showing immune dysregulation and delayed response to infection in older rhesus macaques ([49,52,70]; see also Supplementary Methods). To examine how immunosenescence may impact how changes in social centrality with age influence accumulated disease costs, we included parameters that allowed us to increase susceptibility to infection (likelihood of acquiring infection; parameter: *ai*), duration of infection (parameter: *adi*) and severity of infection (per timestep cost of infection; parameter: *aci*) linearly with age. We simulated pathogen spread across different combinations of these model parameterisations to provide data linking infection cost to age and social network centrality.

#### Model details

We used a stochastic, discrete time implementation of the SIS model. Individuals could either be susceptible (S) or infected (I). (The model code also includes a recovered state (R) but we set its duration to 0 for this analysis so that individuals transitioned immediately from infected back to susceptible). Individuals stayed in a susceptible state until they interacted with an infected individual. For each interaction with an infected individual, a susceptible individual had a probability of transitioning into the infected state of

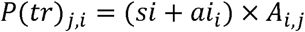

where *P*(*tr*)*_j_*_,*i*_ is the probability of transmission from individual *j* to individual *i*, *si* is a baseline transmission probability, *ai_i_* is the age effect on susceptibility to infection and *A_i_*_,*j*_ is the (undirected) edge weight between *i* and *j* in the grooming network. The *ai_i_* term allowed us to either assume that the probability of infection was homogeneous for every individual (when set to a constant value) or that the probability of infection changed approximately linearly with age so that immunosenescent individuals were more susceptible to infection. *A_i_*_,*j*_ allowed the probability of transmission to depend on how often pairs of animals groomed one another, with this relationship assumed to be linear.

Once an individual is infected, the cost of each infection was calculated as

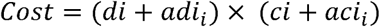

where *di* is the baseline duration of infection (set at 5 time steps for this analysis) before returning to the susceptible (S) state, *adi* is the age effect on infection duration that allows a linear increase in infection duration with age (to a maximum of 15 time steps), *ci* is the baseline per time step cost of infection (set at 1 for this analysis) and *aci* is the age effect on infection severity that allows a linear increase in infection severity per time step with age (to a maximum of 3).

#### Model Parameterisation

We ran in total 24 combinations of parameters (24 models, Table S1). In all versions of the model, we fixed the parameters *di* = 5 (baseline infection duration; considered representative of a typical respiratory infection if a time step is considered to be a day [48,71]; see also Supplementary Methods), *dr* = 0 (baseline duration of recovered period; set to zero to model SIS epidemiological dynamics), *ci* = 1 (arbitrary). We also used a correction such that *A*(*model*)*_i_*_,*j*_ = *A*(*data*)*_i_*_,*j*_ ^0.7^ to facilitate the calculation of the transmission probability per edge by slightly increasing the importance of weak connections for transmission dynamics. We varied the parameters *si* (baseline transmission probability), *ai* (age-based susceptibility to infection), *adi* (age-based duration of infection) and *aci* (age-based severity of infection).

- We varied *si* to have low (0.45), medium (0.6) and high (0.75) transmissibility values. These values were selected so that equilibrium pathogen prevalence equated to a basic reproductive ratio (R_0_) of approximately 1-1.5, 1.5-2 and 2-3 respectively (the R_0_ varied considerably between groups). R_0_ was estimated using the approximation R_0_ = 1/(1-prevalence), where prevalence corresponded to the proportion of infected individuals in a group.
- We varied *ai*, *adi* and *aci* to either be independent of age (no age effect) or increase linearly with age (age effect). *adi* had a maximum value of 10 so that the duration of infection was between 5 (for the youngest) and 15 (for the oldest) time steps, and *aci* had a maximum value of 2 so that the per time step cost of infection was between 1 (for the youngest) and 3 (for the oldest). All values of *adi* were rounded to the nearest whole number.

Simulations contained all possible combinations of *ai*, *adi* and *aci* being ‘on’ (linear age effect) or ‘off’ (no age effect) resulting in 8 possible parameterisations for each transmission probability, and 24 parameter combinations in total (see Table S1).

#### Simulations

Each simulation run consisted of applying the epidemiological model to a given draw from a BISoN generated posterior (a weighted network) for 500 time steps. We applied each parametrisation of the model to each network that we generated - 1000 BISoN posterior networks from each of the 23 group-years (23,000 networks x 24 parameterisations) - resulting in 552,000 simulation runs in total. From each simulation run we calculated the total cost of infection across the whole time period for each individual. Total individual infection costs were collated together with individual data and measures of social network centrality (*see Social Network Construction*) for subsequent analyses.

### Analysis

We used linear mixed effects models (Gaussian error distribution) to summarise the outputs from our epidemiological models (nb. we focus on effect size estimates rather than p-values here as our aim is description rather than inference). We fitted three different linear mixed effects models to understand the relationship between age and sociality on infection cost. For all models we used the same random effect structure, fitting group-year, individual ID and combined group-year and simulation number as random intercepts. We first fitted a model with age as the only continuous fixed effect (model 1) to assess how infection cost changed with age under different model parameterisations (i.e., combinations of transmission probabilities and immunosenescence). We then fitted a model with age, degree, strength, and closeness as fixed effects (model 2) to assess whether the effect of age on infection cost was in part being buffered by social variation across ages. We standardized (z-scored) these three measures of social centrality within group-year and included them as categorical variables in the model to improve fit. We defined 5 categories for each social centrality measure (degree, strength and closeness) as follows: < −1.5 = very low, −1.5 to −0.5 = low; −0.5 to 0.5 = average; 0.5 to 1.5 = high; >1.5 = very high. The estimate of age in model 1 represents the effect of age on infection cost that incorporates any effects of age-related variation in social centrality measures. The estimate of age in model 2 represents the effect of age that occurs independently of the effects of degree, strength, and closeness, and therefore represents the effect of age on infection cost in the “absence” of social effects. This approach allowed us to calculate the overall protective effect of social ageing on age-based infection cost by subtracting the age estimate in model 2 from the age estimate in model 1. If this difference was negative, it would indicate that age-based infection costs were lower when the social effects were not controlled for, indicating that age-based variation in social centrality buffers the effects of age on infection cost and therefore has a protective effect. Finally, we fitted a linear mixed effects model that included an interaction between age (continuous) and each of the social centrality metrics (categorial) to better understand how each of the different social metrics contributed to this overall protective effect (model 3). Fitting an interaction term with each social metric allowed us to assess how the effects of degree, strength, and closeness on infection cost varied with age.

Our first prediction (see Introduction) was that being less socially central at old ages would be more beneficial in terms of reducing infection cost than at young ages. To test this idea, we used the results from model 3 to calculate the reduction in infection cost that resulted from having “average” rather than “high” social centrality when old (18 years old) versus when young (8 years old). We chose these ages for our “young” and “old” categories as age 8 reflects the lower quartile of data and age 18 is the median age of death for females that survive to reproductive age in this population [20,47]. By including all three social metrics in model 3 we could also determine their relative effect on infection risk and therefore assess prediction 2 - that most of the decrease in infection cost associated with social ageing would be driven by older individuals having lower degree and closeness. Finally, by applying models 1-3 across all parameter combinations, we tested prediction 3 - that these protective effects of social ageing would be stronger under conditions where there was greater immunosenescence with age. All analyses were conducted in R version 4.3.1 and modelling was conducted in R version 4.1.1.

## Results

### Cost of infection increases with increased social centrality

On average, there were 51 females in each network (range: 19-73 females) and 169 grooming bouts per network (range: 33-392 bouts). Average network degree, strength and closeness centrality were 9.60 (range: 5.46 - 14.26), 0.62 (range: 0.40 - 0.88), 0.0008 (range: 0.0003 - 0.0025), respectively. All social centrality metrics were weakly to moderately correlated (see Table S2). Each of the three social centrality measures had independent, positive effects on infection cost under all parameter combinations (Fig. 1B,D,F). The effects of strength and closeness on disease cost were particularly strong (Fig. 1D,F), while the effects of degree were more moderate (Fig. 1B). These results confirmed that more socially central individuals suffer greater costs of infection.

**Figure 1.**
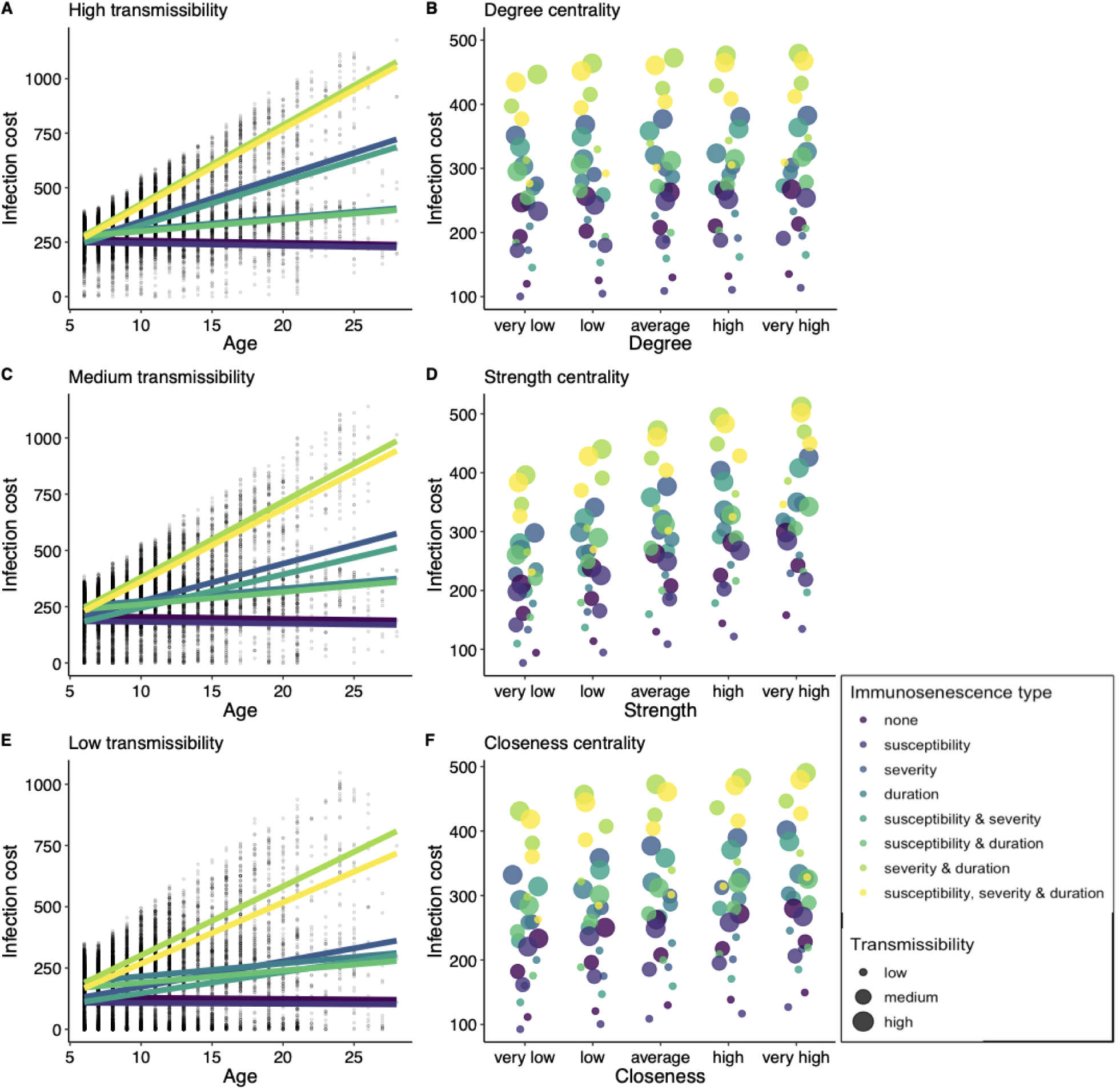
Predicted age and social centrality effects on accumulated infectious disease costs in our simulation results. Effect of age on infection cost at A) high transmissibility, C) medium transmissibility and E) low transmissibility. Across all transmissibilities, infection costs decrease slightly with age with no immunosenescence or immunosenescence only in infection susceptibility, but otherwise increase with age. Points in panel A, C, E represent a random sample of simulation data. Infection costs are higher for individuals with B) higher degree, D) higher strength and F) higher closeness. Transmission probability is represented by point size in B, D, F. In all panels, colour represents the combination of immunosenescent effects included.

### In the absence of immunosenescence, lower infection cost in old age is mediated through lower social connectedness

Under conditions with no immunosenescence (no change in susceptibility (*ai*), severity (*aci*), or duration (*adi*) of infection with age), age had a negative effect on infection cost at all transmissibilities (high: β = −1.09 +/- 0.25; med: β = −0.86 +/- 0.23; low: β = −0.44 +/- 0.17; Fig. 1A,C,E). This means that, for example, at high transmissibility, each year increase in age results in a decrease in 1 cost unit, equating to a reduction of 22 cost units or 4.4 infections between the ages of 6 (our minimum age) and 28 (our maximum age). At the population level, all three sociality measures were negatively correlated with age in our data (degree: r = −0.11; strength: r = −0.13; closeness: r = −0.12; all p < 0.001; Fig. S2). When we included these social centrality measures in the model with age (model 2), the age estimate became less negative (high: β = −0.26 +/- 0.06; med: β = −0.07 +/- 0.05; low: β = 0.20 +/- 0.07) compared to the age estimate in a model with just age (model 1). This indicates that most of the variation that was explained by age in model 1, is now explained by sociality in model 2, suggesting that lower social centrality among older individuals is responsible for a substantial part of the reduction in infection cost with age. For example, at high transmissibility, the protective effect of our three measures of social centrality (i.e., the difference between the age estimate in models 1 and 2) was estimated to be −0.83 (−1.09 - −0.26), meaning that age-based variation in strength, closeness and degree were expected to account for a reduction in 0.83 cost units per year of age, which translates to a reduction of about 3.7 infections between individuals of the minimum and maximum ages in our study (Fig. 2A). We found that this protective effect of social ageing was greatest at medium and high transmissibility and somewhat less prominent at low transmissibility (Fig. 2A).

**Figure 2.**
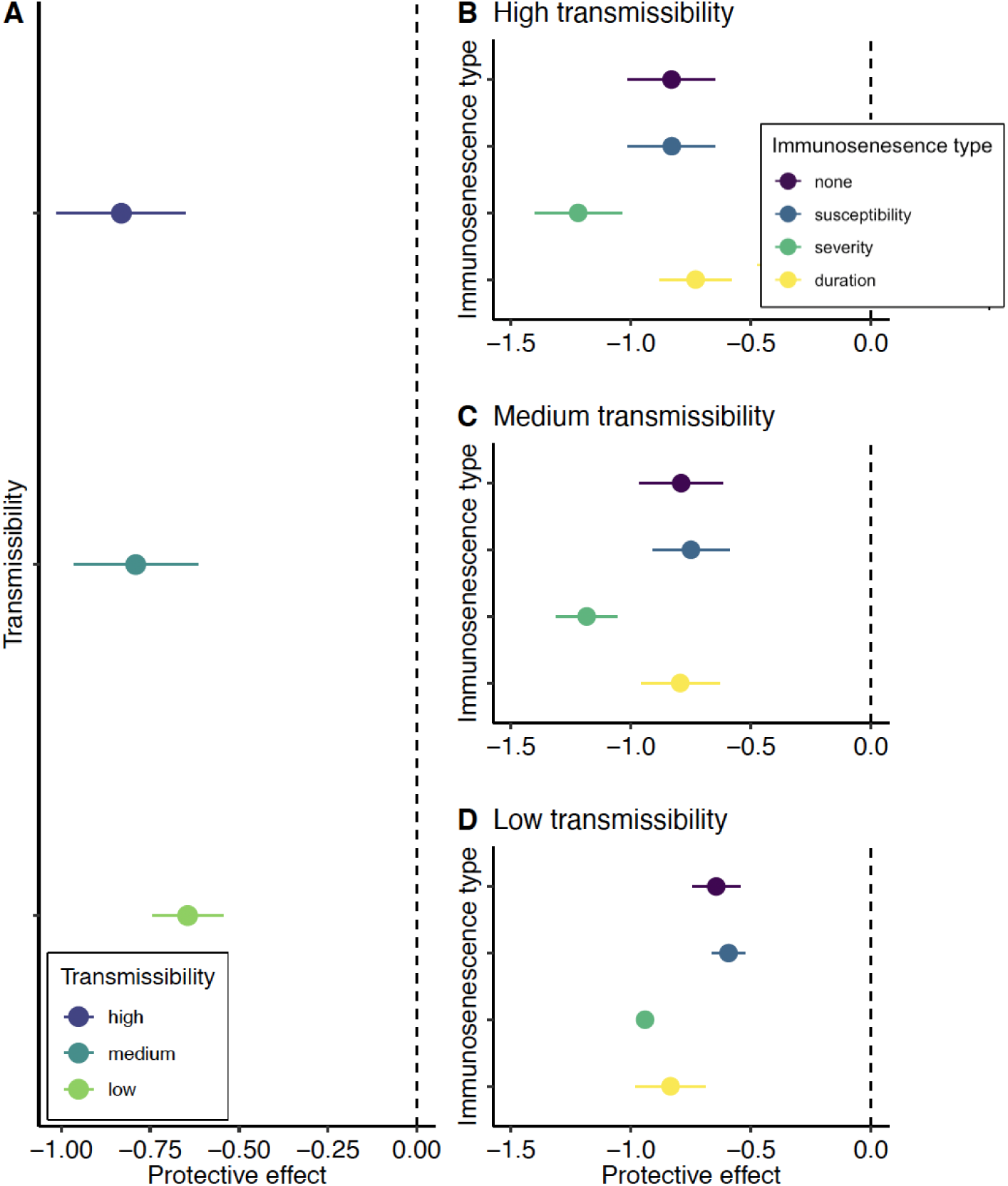
The protective effect of social ageing, measured as the difference in the age estimate between model 1 (without social covariates) and model 2 (with social covariates). A more negative protective effect indicates that the effect of age on infection cost was lower when social centrality measures were not included in the model (model 1), compared to when they were included (model 2). Therefore, the stronger the negative effect, the more variation in infection cost was explained by age-related differences in sociality. Results show the protective effect of social ageing: A) across different transmission probabilities when there is no immunosenescence, and B-D) across each (independent) form of immunosenescence (none, susceptibility, severity, duration) at each transmission probability (high, medium, low).

### The protective effects of lower social connectedness in old age depend on epidemiological characteristics

As expected, linear increases in susceptibility, severity, and duration of infection with age led to infection cost increasing with age, although the strength of this effect depended on the specific combination of parameters in the model (Fig. 1A,C,E). The one exception was when there was only an increase in susceptibility with age, changes in infection cost were similar to those when there was no immunosenescence (Fig. 1A,C,E). In line with prediction 3, the protective effects associated with social ageing tended to get stronger when there was immunosenescence, but this was not always true as it depended on the type of immunosenescence (e.g., whether there were changes in susceptibility, severity, or duration of infection with age) as well as the transmissibility of the pathogen (Fig. 2 & Fig. S3). The protective effects of social ageing were greatest under conditions where there was an increase in infection severity with age, across transmissibilities (Fig. 2B-D). For example, under high transmissibility, when there was an increase in severity with age, the estimate for age in model 1 (just age) was 20.9 while the estimate for age in model 2 (age + sociality) was 22.1. The protective effect was therefore −1.2 meaning that social ageing reduced infection cost by 1.2 units for each year of increase in age. When there were age-based increases in susceptibility, on the other hand, age-associated differences in sociality provided no additional protective effects relative to baseline (i.e., no immunosenescence; Fig. 2B-D). When there were changes in duration of infection with age, the protective effect of social ageing was more complex and depended on the pathogen transmissibility. Relative to when there was no immunosenescence, when transmissibility was low and duration of infection increased with age, social ageing provided more of a protective effect. However, when transmissibility was medium or high and duration increased with age, social ageing provided no additional protective effect compared to baseline. Our results therefore suggest that age-based changes in sociality are likely to provide the greatest benefit when there are increases in the severity or duration of infection with age, or a combination thereof (for a full breakdown of the protective effects across all parameter combinations see Fig. S3).

### Old individuals benefit most from having lower strength and closeness

When considering interactions between age and each of the social metrics (model 3), the reduction in infection cost associated with lower social centrality in old age came primarily from having lower strength and closeness, which ran somewhat contrary to our expectations (see prediction 2 above). Strength and closeness showed clear sociality-age interactions whereby having “average” strength rather than “high” strength when old (18 years) resulted in a substantially greater decrease in infection cost than having “average” strength rather than “high” strength when young (8 years) (Fig. 3). For example, under high pathogen transmissibility and increased infection susceptibility and severity with age, having average rather than high strength when young decreased infection cost by 21.6 units (equivalent to 4.3 infections for a young individual), while having average rather than high strength when old decreased infection cost by almost double this amount (39.0 units; 7.8 infections for a young individual) (Fig. 3B,H).

**Figure 3.**
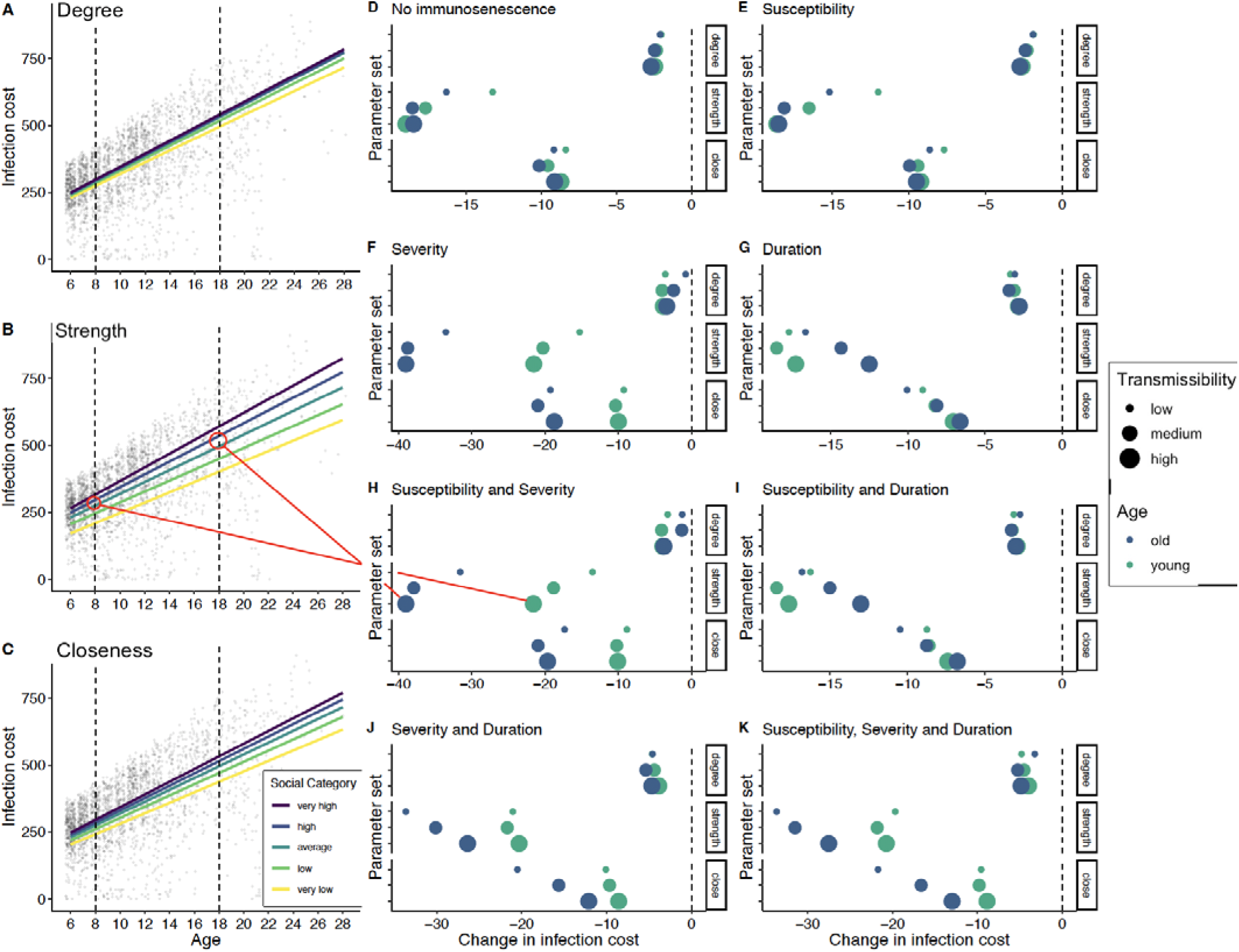
An illustration of how social network centrality and age interact to determine accumulated infection costs. A-C) Predicted age effects on infection costs separated by social centrality category for A) degree, B) strength and C) closeness centrality. Results shown for a specific parameter combination where pathogen transmissibility is high and infection susceptibility and severity show linear increases with age. Points represent a random sample of simulation data. D-K) The change in infection cost for an individual moving from high social centrality to average social centrality when old (18 years; blue points) versus when young (8 years old; green points), for each social centrality measure. Different panels (labelled) show results for different combinations of immunosenescence and point size represents transmission probability. Similar results for a change from average to low centrality are shown in the Supplementary Materials.

Similarly, having lower closeness when young reduced infection cost by 10.0 units (2 infections), while having lower closeness when old reduced infection cost by 19.6 units (3.9 infections) (Fig 3C,H). This supported our prediction that lower connectedness would be more beneficial at old than young ages (see prediction 1 above). However, contrary to our second prediction, lower degree had little to no protective effect, meaning that having “average” degree instead of “high” degree was no more beneficial when old than when young. Following on from the example above, lower degree when young decreased infection cost by 3.9 units and when old by 3.8 units (Fig. 3A,H). In fact, in some cases having fewer social partners for a given strength and closeness centrality seemed to reduce infection cost more at young ages than at old ages (Fig. 3F,H). In line with the results described above, the reduction in infection cost associated with having lower strength or closeness when older was dependent on pathogen transmissibility and on the type of immunosenescence. The reduction in infection cost at old ages compared to young ages was most pronounced when there were age-based increases in infection severity (Fig. 3F,H,J,K). This reduction was somewhat dampened when there were also changes in duration of infection with age (Fig. 3J,K), especially at high/medium transmissibility. When there were changes in infection duration with age but no changes in infection severity, lower values of closeness at old ages were no more beneficial than at young ages, except when pathogen transmissibility was low (Fig. 3G,I). Under this parameter combination, having lower strength was actually more beneficial at young ages than at old ages (Fig. 3G,I). Generally, these findings align well with the results described above, suggesting that having lower values of social centrality when old is less protective when there are increases in infection duration with age and transmissibility is high. We explored the robustness of these patterns by also looking at the difference in infection cost when an individual had “average” social centrality versus “low” social centrality and comparing this at young and old ages. We found the patterns highly comparable to those described above (see Fig. S4).

## Discussion

Our model results suggest that social ageing in rhesus macaques (i.e., lower social centrality in older individuals) is associated with reduced costs accrued from socially transmitted infections. We found that infectious disease cost was strongly positively associated with the network centrality of an individual, as would be expected. Specifically, we found that strength and closeness centrality had strong, independent effects on the overall cost of infection, while the effects of degree on infection cost were more moderate. Given the probability of infection in our model depends on edge weight, the importance of strength on overall cost of infection was expected but nevertheless reflects the widespread importance of strength in explaining infection risk in free-living populations [30]. Under conditions with no immunosenescence (no increases in infection susceptibility, severity, or duration with age) older individuals accrued a lower cost of infection compared to younger individuals across all transmissibilities modelled, with the benefits peaking for pathogens with intermediate-high transmission probabilities. These lower costs of infection were driven by age-associated differences in social centrality, as all three social centrality metrics were negatively correlated with age and including centrality measures within the same model resulted in the protective effect largely disappearing. When we included (linear) immunosenescence we were able to show that having lower social centrality had a much greater benefit for older than younger individuals, with this effect strongest when the severity of infection increased with age.

### The protective effect of lower social centrality on infection risk in older individuals

We found that older individuals accrued lower disease costs than younger individuals in the absence of immunosenescence. Given previously documented within-individual declines in closeness centrality and degree with age [21,25], we anticipated that this protective effect would be driven by age-based differences in social centrality, which our results support. Our analyses, however, were not explicitly longitudinal, meaning that we cannot fully rule out selective disappearance of more central individuals partly explaining these results. It is possible that the high infectious disease risk associated with being more socially central leads these individuals to die earlier. However, given that social centrality is associated with survival in this population [8,20,42], it seems likely that longitudinal changes in social network position reduce older individual’s exposure to infection. This is likely to be particularly true for degree and closeness for which we have clear evidence that population level differences are driven by within-individual changes with age [21,25]. The fact that strength was negatively correlated with age and contributed substantially to the protective effect provided by social ageing was surprising given that previous analyses in this study system have found a non-significant (although weakly negative) relationship between age and strength both within and between individuals [21]. However, it is possible that, combined, this weakly negative relationship at both the within and between-individual level results in a more substantial population-level decline in strength with age, which we detect in our analyses. It is also possible that the inclusion of more data than previous analyses has enabled us to detect change in strength with age not previously evident.

Overall, we found the protective effect of social ageing to be strongest for pathogens with intermediate-high transmissibility and weaker for less transmissible pathogens. That is, the reduction in infection costs with age (in the absence of immunosenescence) was highest when transmissibility was high and pathogen prevalence (i.e., proportion of infected individuals in the group) within groups was therefore relatively high (>30% or more), with a weaker reduction when transmissibility, and therefore pathogen prevalence, in groups was low. There are a variety of pathogens that are transmitted via direct contact, and which vary in their level of transmissibility, which could be represented by our model. For example, Shigella (a bacterial pathogen) is transmissible by close contact in macaques [62] and has spread very rapidly through the Cayo Santiago macaque population previously [72]. Additionally, macaques are known to be infected by a range of respiratory viruses (e.g, coronaviruses, influenza viruses), which will likely fall across the range of transmissibilities considered in our study (see Supplementary Methods), suggesting our findings are likely to be highly relevant for real-world pathogens.

### Immunosenescence enhances the protective effect of lower social centrality in older individuals

When adding immunosenescence to our models, we found that the protective effects of social ageing were generally stronger than under conditions with no immunosenescence. Overall, we found that lower levels of social centrality at old ages led to the most substantial reductions in infection costs when infection severity increased with age. If we instead considered conditions where infection duration increased with age, then the protective effects of social ageing depended on pathogen transmissibility. That is, lower connectivity at older ages was more important for decreasing disease costs at low compared to high transmissibility when infection duration increased with age. These results can be explained intuitively based on the epidemiological dynamics of high versus low pathogen transmissibility within an SIS model. When transmissibility, and therefore (in our model) prevalence is very high, individuals will typically be re-infected very quickly once they recover [73], meaning that the duration of each infection is less important to overall disease cost. However, when transmissibility, and therefore prevalence, is low it may take a while for an individual to be re-infected once it recovers from infection. As a result of this reduced frequency of infection, the fact that each infection lasts longer is disproportionately costly. Therefore, having lower social centrality when old relative to when young under this low transmissibility scenario has a more substantial protective effect. When we looked at conditions where infection susceptibility increased with age, we found that older individuals gained very little protective effect from having lower social connectivity relative to younger individuals.

It should be noted that comparisons of the relative importance of age-related changes in susceptibility, severity and duration for disease cost are limited somewhat by how their relative effects match (e.g., infection severity can be 3 times higher for our oldest compared with our youngest individual, while the difference caused by changes in susceptibility is between 1.33 and 1.56 times depending on the baseline transmission probability). These differences arise because parameters were chosen to be reflective of a reasonable parameter space for many respiratory infections rather than specifically for comparisons of relative effects. Despite this limitation, we can still draw some general conclusions. When pathogen transmissibility is high, most of the age-based differences in accumulated disease cost are driven by increases in the severity of infection with age, while increases in infection susceptibility or duration with age increase age-based infection cost only minimally. When pathogen transmissibility is lower, age-related changes in infection duration become relatively more important for age-based infection costs, although changes to severity still dominate. Generally, this suggests that age-based differences in social centrality are most likely when there are increases in severity of infection with age. In the case of a pathogen with low transmissibility, increases in duration of infection with age might also result in social declines.

### Social trade-offs in old age and the role of infectious disease

The measures of social centrality that were most important for cost of infection in older individuals were not exclusively those known to change within-individuals as they age, indicating that there may be potentially important social trade-offs. We found that, in general, older individuals would benefit most from having lower strength followed by lower closeness centrality, with lower values of degree having smaller and less consistent effects on infection risk. When we contrast this with observed within-individual changes in social centrality [21,25], it is noteworthy that individuals in this population show behavioural changes with age that reduce both their degree and closeness centrality but not their strength. Although here we have shown that strength is negatively correlated with age it remains unclear whether this negative relationship is driven by within-individual changes or between-individual differences because of selective disappearance or differences between cohorts. Strength to top partners has a positive link to health and survival in rhesus macaques [42] and diverse group-living species [1]. Therefore, it seems possible that while having lower strength may be the most effective way for older animals to cut infectious disease risk, the costs of doing so may be high, and preserving strong social relationships may itself be an important form of social buffering that can protect against infectious disease ([74]; see Limitations section below). By maintaining strong social connections and avoiding interactions with less familiar or new social partners [21,25] individuals can instead reduce other aspects of their social centrality (e.g., closeness) that also reduce infectious disease costs but which are less strongly associated with other aspects of health and fitness. In this way, older individuals may still reap the benefits of social relationships while minimizing the risks of infection [35].

While, to date, most research on social ageing has focused on age-based differences in direct connectedness (cf. [23,25,26]), measures of “flow” through a network, as captured through indirect metrics, can also be highly relevant for pathogen transmission [30]. Our previous work has shown that by changing simple behavioural rules (in this case, reassociating with the same partners and mixing less widely with the broader network) ageing individuals can facilitate changes in both their direct connectedness (i.e. degree) and their indirect connectedness (i.e. closeness) [25]. Here we have seen that it is the effects of these age-based behavioural changes on closeness which are particularly beneficial, relative to the effects on degree, for mitigating infection risk. Furthering our understanding of the intersection between social ageing and infectious disease therefore necessitates deepening our understanding of how behavioural changes with age facilitate changes not only in direct connectedness but also changes in connectedness to the wider network.

### Potential implications for the evolution of social ageing

While our results suggest that individuals could benefit from reduced infectious disease risk by reducing their social networks in old age [21,25], more work is required before we can say that social ageing is adaptive. While, it is appealing to consider that social ageing might be an evolved strategy to counteract declines in immunity and associated increases in disease burden in later life, the strength of selection on a trait will typically decline with age [75,76]. This has two implications for the evolution of social ageing in response to infectious disease risk. First, it means that when social network centrality influences fitness we would expect social behaviour to senesce as a result of weakening selection in later life [75], independent of any late-life benefits of reduced social centrality. This expected decline could even be enhanced in rhesus macaques where positive effects of social relationships on survival are reduced for older individuals [20]. Second, under most conditions the declining strength of selection with age makes it less likely that social ageing could have evolved to reduce the costs attributed to infection late in life when immunosenescence increases the risk and severity of infections [35]. However, substantial immunosenescence at ages where individuals still have some reproductive value could cause infectious disease to influence social ageing. It may be in these cases that observed patterns of social ageing are a plastic response that can be selected for to mitigate rapid declines in immune performance with age. Additionally, predictions related to the strength and direction of demographic selection at old ages can depend on assumptions about density-dependent population regulation [77–79], and it is not immediately clear where mortality related to infectious disease fits in this context. Therefore, any selection acting on age-related changes in social behaviour are likely to be highly context dependent and may change depending on which aspects of sociality predict fitness earlier in life too. Expanding theory to integrate age-related changes in social behaviour within existing evolutionary models of age-dependent mortality and senescence will be key to better understanding the role of infectious disease in social ageing.

### Limitations

There are some limitations to our model that should be considered when interpreting our results and that offer fruitful choices for future work. First, we have considered only directly transmitted pathogens and focused on SIS (susceptible-infected-susceptible) epidemiological dynamics. These methodological choices make sense. Directly transmitted pathogens are widespread threats to human and non-human animal health [80] and their epidemiological dynamics will be most strongly influenced by social interaction patterns [81]. However, incorporating indirectly transmitted pathogens may be important if co-infection affects morbidity [82,83]. We focused on SIS dynamics as this provides a convenient way to model endemic disease without explicitly incorporating demography or immune dynamics. Future work could build on ours by testing how the choice of disease model can affect results, or by developing long-term integrated network-demographic models of disease.

We have also assumed a linear relationship between interaction duration and the probability of infection. However, for some pathogens with high transmissibility almost any social contact may be sufficient, while for others only prolonged interactions may allow the pathogen to spread. These differences would likely mean that the impact of social ageing on pathogen transmission will depend on the shape of the dose-response curve. For example, we might predict that social ageing is ineffective against mitigating disease risks for pathogens that can spread easily via even short duration proximity. While social relationships are known to expose individuals to disease risk, they can simultaneously help individuals cope with infections once acquired [74,84]. We have not explicitly modeled these ‘social support effects’, which we may expect to play a role in macaques [74]. In general, we would expect the greatest social support effects from an individual’s strongest social relationships [42,85,86], which may amplify any protective effects of social ageing against disease as long these types of relationships are maintained [21,27]. Existing models have incorporated social support effects into network models of infectious disease spread [87], and using them explicitly in this context could provide additional insight.

Further, while we have focused here on how differences in social behaviour can affect infection risk we have not considered how the spread of infection can also influence social behaviour and network structure. In some cases, sickness behaviour and social avoidance of infected individuals can change an individual’s social interactions (reviewed in [88,89]) influencing the structure of the network [90]. Given older individuals tend to be more susceptible to pathogens, lower connectedness with age could be a consequence of, rather than a proactive response to, infectious disease risk. However, this seems an unlikely explanation of social ageing in our system given evidence that older females appear to be more selective in their partner choice rather than arbitrarily decreasing their connectedness as might be expected with sickness behaviour or being avoided by other individuals [69]). Alternatively, if social selectivity is selected for by infectious disease costs (see caveats above), then older individuals might actually be hyper-responsive to cues of infection, leading to greater infection avoidance behaviour with age. Older individuals’ may also be better able to detect infections in the small number of individuals they know well and this could enhance avoidance strategies [35]. Ultimately, how social ageing fits into the co-dynamics of social behaviour and infectious disease spread is likely to be complex and warrants further research.

### Conclusions

We used epidemiological models to demonstrate that reduced social connectedness in old age has the potential to provide a protective effect against the accrued costs of endemic infectious diseases in a free-ranging population of rhesus macaques. By considering the impacts of different forms of immunosenescence we showed that the benefits of social ageing could vary considerably depending on the interaction between pathogen traits (transmissibility) and how changes to an individual’s immune performance manifest in terms of susceptibility, severity and duration of infection. In addition, the aspects of social centrality that most impacted disease costs for older individuals were not exclusively those that we know change within-individuals as they age, highlighting the trade-offs inherent to interacting with others. Although we focused in this paper on how age-based changes in susceptibility to infectious disease might facilitate social ageing, the other way that immunosenescence might affect age-based changes in sociality is through reduced healing ability. Being less able to recover from wounds might impose greater social costs at old ages leading individuals to alter their social centrality to avoid competitive interactions [12]. Generally, understanding how immunosenescence intersects with age-based variation in sociality remains an open and intriguing question. Our results demonstrate the clear potential for infectious disease to influence social ageing and point towards the value of developing new theoretical models that consider the evolutionary dynamics involved.

## Acknowledgements

Thank you to J. Firth, G. Albery, S. Bouwhuis and R. Salguero-Gomez for organizing this special issue. We are grateful to the Caribbean Primate Research Center for maintaining the Cayo Santiago population and for access to the study site, and all the field technicians who have contributed to the long-term behavioural database over the years. Thank you to S. Ellis, D. Nussey, and J. Moorad for thoughtful discussion during the manuscript’s development. This work was supported by the following grants from the National Institute of Health (NIH): grant nos R01-AG060931, R00-AG051764, R01-MH096875, R37-MH109728, R01-MH108627, R01-MH118203, U01MH121260, R01-NS123054 and the Kaufman Foundation: grant no KA2019-105548. The Cayo Santiago Field Station is supported by the Office of Research Infrastructure Programs of the NIH (2P40OD012217). MJS is supported by a Royal Society University Research Fellowship URF\R1\221800.

