## Supplementary Materials for "Social ageing can protect against infectious disease in a group-living primate"

**This PDF file includes:**

Supplementary Methods
Tables S1 to S2
Figures S1 to S4
SI References

**Supplementary Methods**

We specified our susceptible-infected-susceptible (SIS) epidemiological model to broadly reflect typical respiratory pathogen spread by close contact.

As detailed in the main text, we selected three values for our transmission rate parameter (*si*) such that created distinct stable endemic prevalences across our 23 group-years. These stable prevalences equated to approximate R_0_ values of 1-1.5, 1.5-2 and 2-3 for the low, medium and high transmissibility conditions respectively. This compares, for example, to R_0_ values of influenza viruses (R_0_=1.2-1.7; [[1]](https://paperpile.com/c/qsFCBs/kcQF)) and SARS-CoV-2 (original variant R_0_=2.5; [[2]](https://paperpile.com/c/qsFCBs/ikDY)) in humans (where the R_0_s of respiratory pathogens are better characterised).

The minimum duration of infection (*di*) in our model was 5 days, increasing to a maximum of 15 days in our oldest individuals when there was immunosenescence acting on infection duration (*adi*). In the immunosenescent treatment most individuals will have an infection duration intermediate between these values but closer to the minimum value. These values are broadly representative of the infection duration of respiratory viruses in laboratory macaques. For example, rhesus macaques infected intranasally with SARS-CoV-2 test positive for viral RNA in throat swabs 3-7 days and nasal swabs for 6-9 days [[3]](https://paperpile.com/c/qsFCBs/SVbm), while influenza viruses show a peak viral load that lasts approximately 5-9 days in rhesus macaques [[4]](https://paperpile.com/c/qsFCBs/VJyb) and 4-9 days in closely-related long-tailed macaques [[4]](https://paperpile.com/c/qsFCBs/VJyb). While clinical scores can occasionally last longer (e.g. [[5]](https://paperpile.com/c/qsFCBs/1kaR)), they also generally drop after 8-9 days in healthy individuals (e.g. [[6]](https://paperpile.com/c/qsFCBs/kPch)) and we need to find the correct balance between accounting for an individual being infectious and suffering from symptoms. There is evidence for extended duration of infection by SARS-CoV-2 in older rhesus macaques (e.g., up to 14 days in 15 year old individuals; [[7]](https://paperpile.com/c/qsFCBs/JoBA)) providing support for our extended infection duration in the immunosenescent treatment.

As described in the main text, our baseline cost of infection (*ci*) was chosen arbitrarily as we were interested in relative differences rather than absolute values. The potential for immunosenescence to lead to increased infection costs in older macaques (*aci*) is supported by various laboratory studies [[4,5]](https://paperpile.com/c/qsFCBs/1kaR+VJyb), with one study indicating clinical scores 2-3x higher in 18 year old vs 3.4 year old macaques [[5]](https://paperpile.com/c/qsFCBs/1kaR), broadly supportive of our maximum potential difference of 3x higher disease costs in (hypothetical) 28 year old vs 5 year old individuals.

| **Parameter set** | **Transmission probability (si)** | **Age based susceptibility (ai)** | **Age based severity (aci)** | **Age based duration (adi)** |
| --- | --- | --- | --- | --- |
| 1 | 0.45 | 0 | 0 | 0 |
| 2 | 0.60 | 0 | 0 | 0 |
| 3 | 0.75 | 0 | 0 | 0 |
| 4 | 0.45 | 1 | 0 | 0 |
| 5 | 0.60 | 1 | 0 | 0 |
| 6 | 0.75 | 1 | 0 | 0 |
| 7 | 0.45 | 0 | 1 | 0 |
| 8 | 0.60 | 0 | 1 | 0 |
| 9 | 0.75 | 0 | 1 | 0 |
| 10 | 0.45 | 1 | 1 | 0 |
| 11 | 0.60 | 1 | 1 | 0 |
| 12 | 0.75 | 1 | 1 | 0 |
| 13 | 0.45 | 0 | 0 | 1 |
| 14 | 0.60 | 0 | 0 | 1 |
| 15 | 0.75 | 0 | 0 | 1 |
| 16 | 0.45 | 1 | 0 | 1 |
| 17 | 0.60 | 1 | 0 | 1 |
| 18 | 0.75 | 1 | 0 | 1 |
| 19 | 0.45 | 0 | 1 | 1 |
| 20 | 0.60 | 0 | 1 | 1 |
| 21 | 0.75 | 0 | 1 | 1 |
| 22 | 0.45 | 1 | 1 | 1 |
| 23 | 0.60 | 1 | 1 | 1 |
| 24 | 0.75 | 1 | 1 | 1 |

**Table S1.** Combination of parameters for our simulations. Transmission probability (si) was set to 0.45 (low), 0.60 (medium) or 0.75 (high). For other immunosenescence parameters (ai, aci, adi); 0 indicates no relationship with age, 1 indicates linear relationship with age.

**Table S2.** Pearson’s correlation coefficients between measures of social centrality.

|  | **Degree** | **Strength** | **Closeness** |
| --- | --- | --- | --- |
| **Degree** | 1 | 0.61 | -0.26 |
| **Strength** | 0.61 | 1 | 0.10 |
| **Closeness** | -0.26 | 0.10 | 1 |

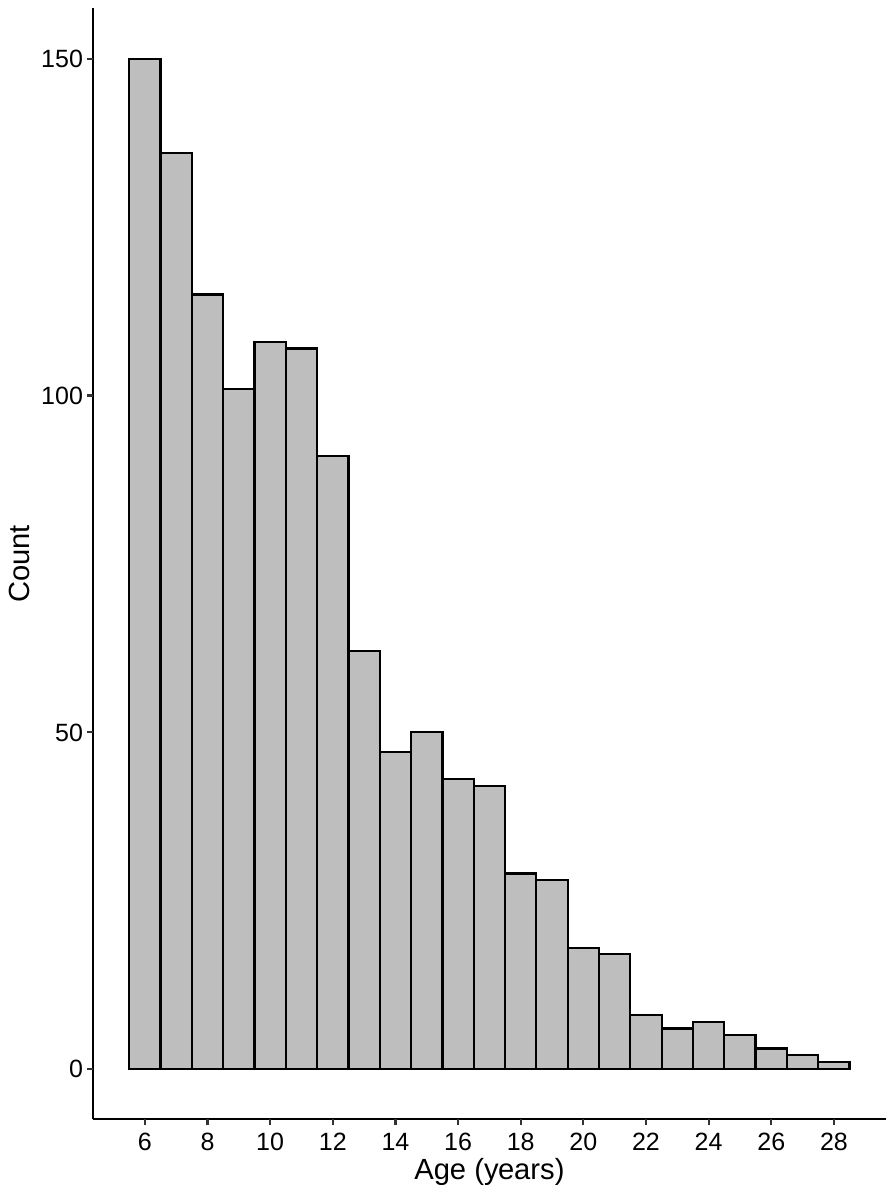

**Figure S1.** Histogram showing distribution of female ages in our population.

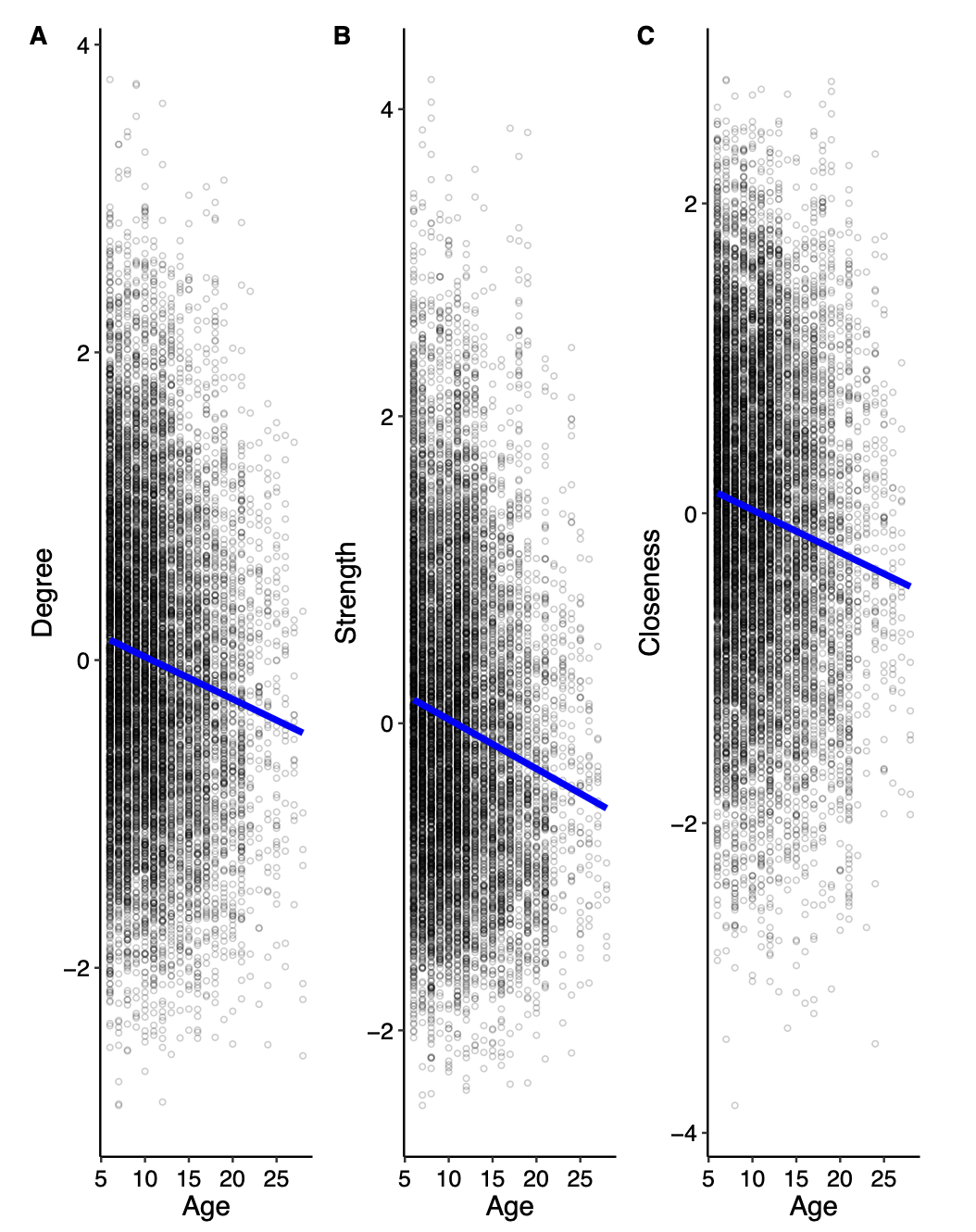

**Figure S2.** Population-level relationship between age and each of the social centrality measures. Social centrality measures are standardized (z-scored) within group-year. Data points represent a random sample of the raw data (1000 networks across 23 group-years) to aid visualisation.

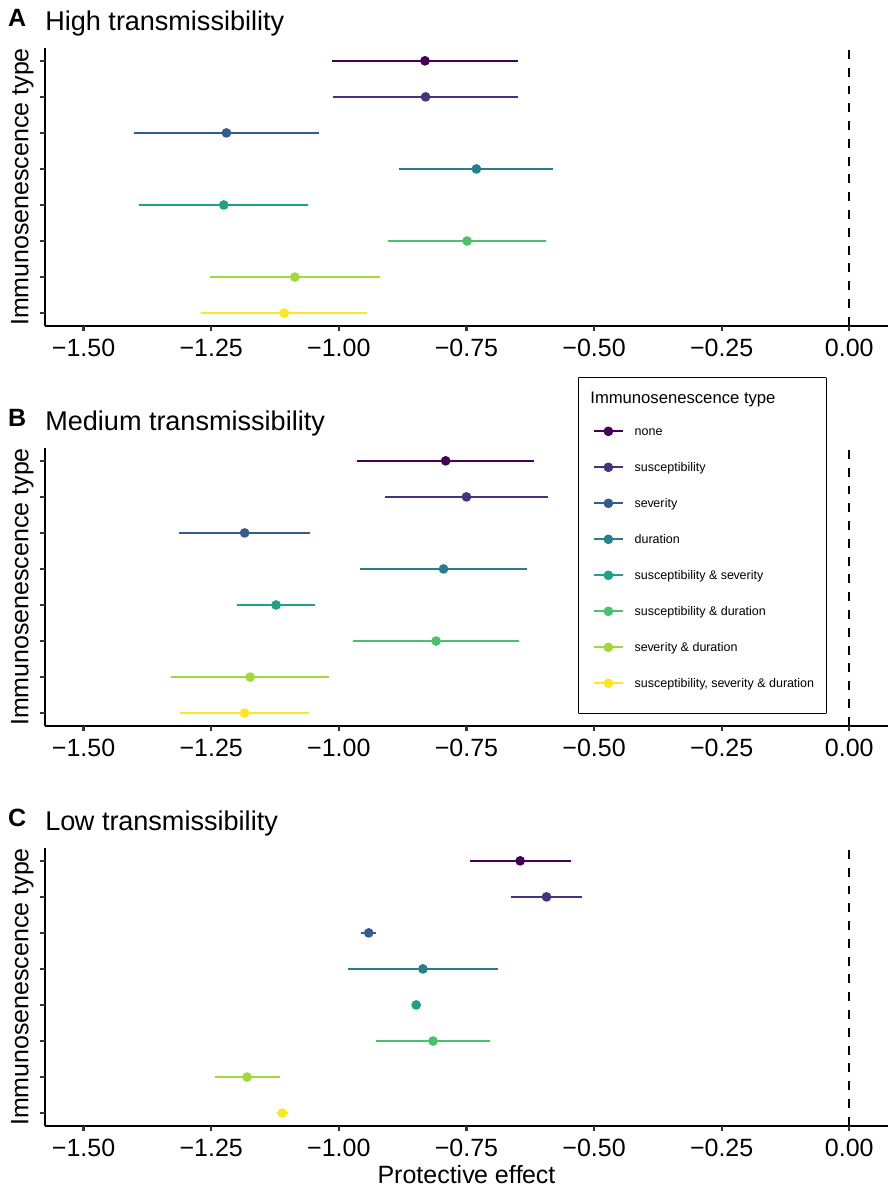

**Figure S3.** The effects of different combinations of immunosenescence on the protective effect of age-associated changes in social centrality against accumulated disease costs for A) high transmissibility, B) medium transmissibility and C) low transmissibility.

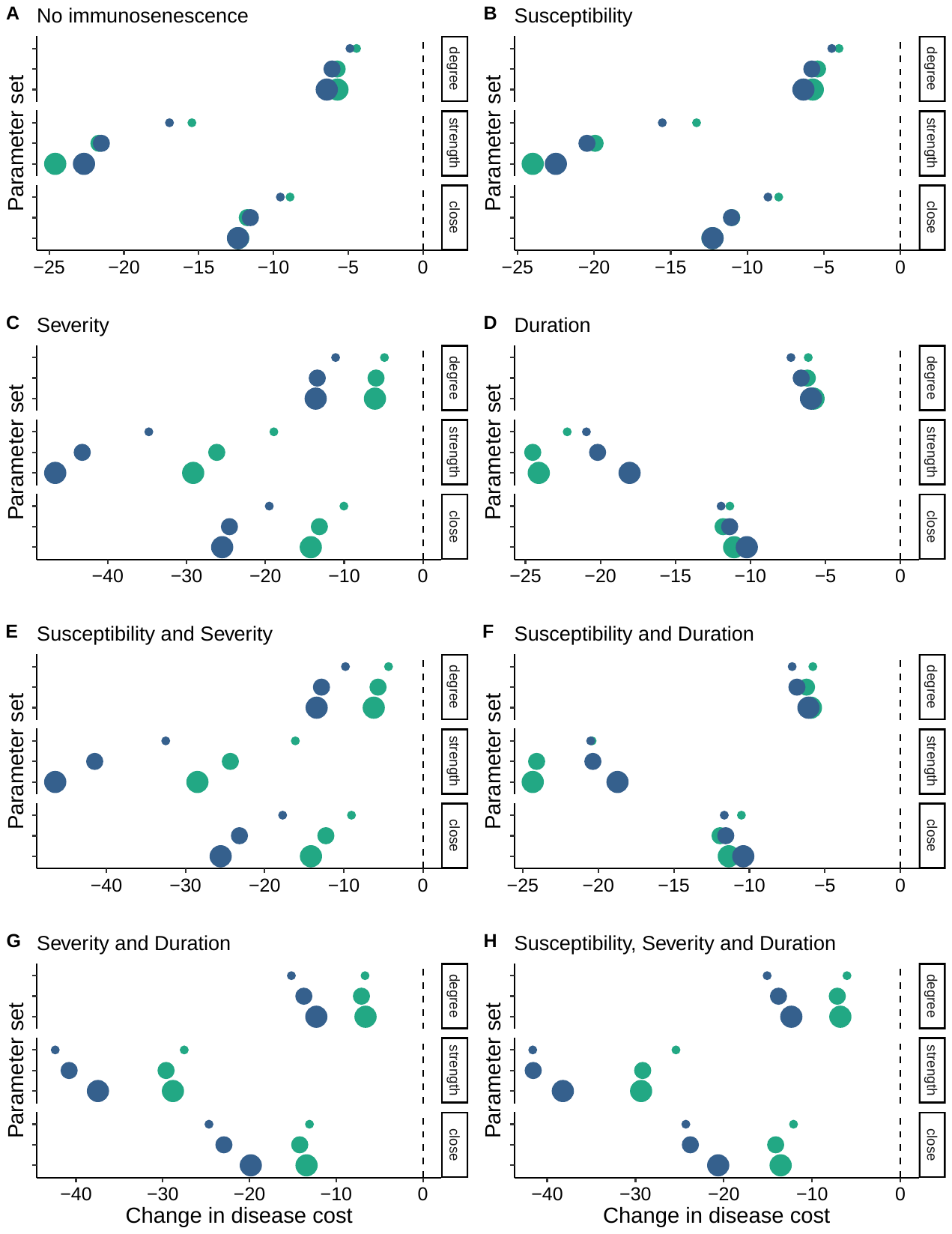

**Figure S4.** The change in infection cost for an individual moving from the average to the low social centrality category for old (18 years; blue points) and young (8 years old; green points) for each social centrality measure. Different panels (labelled) show results for different combinations of immunosenescence and point size represents transmission probability.
